# Identification of potential vaccine candidates against *SARS-CoV-2*, A step forward to fight COVID-19: A Reverse Vaccinology Approach

**DOI:** 10.1101/2020.04.13.039198

**Authors:** Ekta Gupta, Rupesh Kumar Mishra, Ravi Ranjan Kumar Niraj

**Author notes:** **Correspondence Author:** Dr. Ravi Ranjan Kumar Niraj, Assistant Professor, Amity Institute of Biotechnology, Amity University Rajasthan, Jaipur (INDIA)-303002.

## Abstract

The recent Coronavirus Disease 2019 (COVID-19) causes an immense health crisis to global public health. The COVID-19 is the etiologic agent of a recently arose disease caused by the severe acute respiratory syndrome coronavirus 2 *(SARS-CoV-2).* Presently, there is no vaccine available against this emerged viral disease. Therefore, it is indeed a need of the hour to develop an effectual and safe vaccine against this decidedly pandemic disease. In the current study, we collected *SARS-CoV-2* genome which is prominent in India against human host, further more using reverse vaccinology here we claim effective vaccine candidates that can be mile stone in battle against COVID19. This novel study divulged one promising antigenic peptide GVYFASTEK from surface glycoprotein (protein accession no. - QIA98583.1) of *SARS-CoV-2,* which was predicated to be interacted with MHC alleles and showed up to 90% conservancy and high value of antigenicity. Subsequently, the molecular docking and simulation studies were verified molecular interaction of this prime antigenic peptide with the residues of HLA-A*11–01 allele for MHC Class I. After vigorous analysis, this peptide was predicted to be suitable epitope which is capable to induce the strong cell-mediated immune response against the *SARS-CoV-2.* Consequences from the current study could facilitate selecting *SARS-CoV-2* epitopes for vaccine production pipelines in the immediate future. This novel research will certainly pave the way for a fast, reliable and virtuous platform to provide timely countermeasure of this dangerous pandemic disease, COVID-19.

## 1. Introduction

The Coronavirus Disease 2019 (COVID-19) begin in December 2019, like a viral outbreak in Wuhan city of China (Holshue et al., 2020). It gained rapid foothold across the world resulting in WHO declared it as a pandemic (https://www.cnbc.com/2020/03/11/who-declares-the-coronavirus-outbreak-a-global-pandemic.html). As on October, 9 2020 worldwide total 36,799,479 cases and 1,067,537 deaths were reported by WHO(https://www.worldometers.info/coronavirus/).Coronaviruses have long been recognized as crucial veterinary pathogens, causing enteric and respiratory diseases in mammals as well as in birds(Adeoye, Oso, Olaoye, Tijjani, & Adebayo, 2020). The SARS-CoV-2 virus spreads primarily through saliva, droplets, or discharges from the nose of an infected person after coughing or sneezing (Pant, Singh, Ravichandiran, Murty, & Srivastava, 2020). The coronaviruses are enveloped RNA viruses having the largest genome among all RNA (Sinha et al., 2020). As no drug or vaccine is available against SARS-CoV-2 many countries, including India has announced lockdown to keep away from further spread of the virus(Umesh, Kundu, Selvaraj, Singh, & Dubey, 2020).

In this period of time when the continuous transmission of the virus across borders and health burden on the global scale is rapidly increasing, more urgent studies are required and in the absence of effective cures majorly drugs, vaccination or immunization therapy is imperative in order to target whole population. Likewise to move forward vaccine development pipeline, immunoinfomatics tools have been proved crucial(S. Mishra & Sinha, 2009). There is less knowledge about the pathogenesis of the virus; therefore, an immunoinformatics-based approach to investigate the immunogenic epitopes is required(Enayatkhani et al., 2020).

Since the Covid19 has affected almost all of the world’s population, promiscuous epitopes binding to a variety of HLA alleles for larger dissemination is vital. For that, *in silico* approaches will be remarkably useful in helping to develop a cure in as fast manner as possible(D. Mishra, 2020). The antibody generation by activation of B-cell as well as acute viral clearance by T-cells along with virus-specific memory generation by CD8+ T-cells are analogously important to develop immunity against the virus(Enjuanes et al., 2016). The S protein is considered to be highly antigenic and thereby can evoke strong immune responses and generate neutralizing antibodies that can block the attachment of virus to the host cells(Du et al., 2009).

In reverse vaccinology, various tools of *in silico* biology are used to discover the novel antigens by studying the genetic makeup of a pathogen and the genes that could lead to good epitopes are determined. The reverse vaccinology approach provide fast and cost effective vaccine discovery plateform(Ullah, Sarkar, & Islam, 2020). In reverse vaccinology approach a novel antigen is identified using omics analysis of target organism. In-silico as well as reverse vaccinology approach facilitates an easier, time and labor saving process of antigen discovery(Gupta, Gupta, & Niraj, 2019).

Herein, we explored the proteome of SARS-CoV-2, prominent in Indian geographical origin against human host to identify potential antigenic proteins and epitopes that can effectively elicit cellular mediated immune response against COVID-19. This significant research disclosed promising antigenic peptides from surface glycoprotein of *SARS-CoV-2.* The outcomes from this very significant analysis could help selecting *SARS-CoV-2* epitopes for vaccine production pipelines soon.

## 2. Methodology

### 2.1. Selection of strain

The highly virulent strain *SARS-CoV-2* was selected for *in-silico* study. The complete genome of *SARS-CoV-2* is available on the National Center for Biotechnology Information or NCBI(https://www.ncbi.nlm.nih.gov/) with *RefSeq NC_045512.2*

### 2.2. Protein Identification and Retrieval

Twelve viral protein sequences of *SARS-CoV2* against (Host: Human, Country: India) were retrieved from ViPR database(Pickett et al., 2012). These proteins were: Orf10 protein (**QIA98591.1)**, Orf8 protein **(QIA98589.1)**, Orf7a protein **(QIA98588.1)**, Orf6 protein **(QIA98587.1)**, Orf3a protein **(QIA98584.1)**, Membrane glycoprotein **(QIA98586.1)**, Envelope protein **(QIA98585.1)**, Surface glycoprotein **(QIA98583.1)**, Surface glycoprotein **(QHS34546.1)**, Nucleocapsid protein **(QII87776.1)**, Nucleocapsid protein **(QII87775.1)** and Nucleocapsidphosphoprotein **(QIA98590.1)**.

### 2.3. Physicochemical Property Prediction

The ExPASy’s online tool ProtParam (Walker, 2005) was used to predict various physicochemical properties of the selected protein sequences.

### 2.4. Protein Antigenicity

VaxiJen v2.0 (Doytchinova & Flower, 2007) was used to predict antigenicity of proteins. This software uses FASTA file format of amino acid sequences as an input and on the basis of physicochemical properties of proteins, antigenicity is predicted by the tool. The output is denoted according to an antigenic score (Meunier et al., 2016). During analysis, the threshold was kept at 0.4.

### 2.5. B-cell and T-cell Epitope Prediction

The B-cell and T-cell epitopes of the selected surface glycoprotein sequence was predicted via IEDB (The Immune Epitope Database). The IEDB database holds large amount of experimental data on epitopes and antibodies. It allows robust analysis on many epitopes in the context of some tools like: conservation across antigens, population coverage, and clusters with similar sequences (Vita et al., 2019). In order to obtain MHC class-I restricted CD8+ cytotoxic T-lymphocyte (CTL) epitopes of the selected surface glycoprotein sequence, NetMHCpan EL 4.0 prediction method was applied for HLA-A*11–01 allele. For MHC class-II restricted CD4+ helper T-lymphocyte (HTL) epitopes were obtained for HLA DRB1 *04-01 allele using Sturniolo prediction method. Top ten MHC class-I and MHC class-II epitopes were randomly selected on the basis of their percentile scores and antigenicity scores (AS). Five random B-cell lymphocyte epitopes (BCL) were selected based on of their higher length using Bipipered linear epitope prediction method(Ullah et al., 2020).

### 2.6. Antigenicity and Allergenicity of the predicted epitopes

VaxiJen v2.0 was utilized to predict protein antigenicity. During antigenicity analysis, threshold was kept at 0.4. The allergenicity of the selected epitopes was predicted via AllerTOP v2(https://www.ddg-pharmfac.net/AllerTOP/).

### 2.7. Transmembrane Helix and Toxicity Prediction of the predicated epitopes

The transmembrane helix of the selected epitopes was predicted using the TMHMM v2.0 server (http://www.cbs.dtu.dk/services/TMHMM/). The server predicts whether the epitope would be transmembrane, remain inside or outside of the membrane. The toxicity prediction of the selected epitopes was carried out via ToxinPred server https://webs.iiitd.edu.in/raghava/toxinpred/protein.php.

### 2.8. Prediction of Conservancy for the Selected Epitopes

The conservancy analysis of the epitopes was performed via the epitope conservancy analysis tool of IEDB server (Vita et al., 2019). During analysis, the sequence identity threshold was kept at ‘>=50’.

### 2.9. The MHC Alleles; Cluster analysis

Cluster analysis was carried out by MHCcluster 2.0 (Thomsen, Lundegaard, Buus, Lund, & Nielsen, 2013). During cluster analysis, the number of peptides to be included was kept at 50,000, the number of bootstrap calculations were kept at 100. For cluster analysis, the NetMHCpan-2.8 prediction method was used.

### 2.10. Generation of the 3D Structures of the Selected Epitopes

The PEP-FOLD3 online tool was used to predict 3D structures of the selected best epitopes (http://bioserv.rpbs.univ-paris-diderot.fr/services/PEP-FOLD3/). This server is a tool for generating de novo peptide 3-D structure (Lamiable et al., 2016; Shen, Maupetit, Derreumaux, & Tuffery, 2014; Thévenet et al., 2012).

### 2.11. Molecular Docking and Molecular Dynamics Simulation

Pre-docking was carried out by UCSF Chimera (Pettersen et al., 2004). The peptide-protein docking of the selected epitopes was carried out by online docking tool PatchDock(https://bioinfo3d.cs.tau.ac.il/PatchDock/php.php) results of PatchDock were refined and re-scored by FireDock server (http://bioinfo3d.cs.tau.ac.il/FireDock/php.php). Later on, docking was performed by HPEPDOCK server(Zhou, Jin, Li, & Huang, 2018). Docking pose analysis was done by using Ligplot(Wallace, Laskowski, & Thornton, 1995). The molecular simulation was executed with GROMACS 2018.1 package with the force field as Gromos43a1. The protein salvation was performed with SPC water model in a cubic box (10.8 U χ U 10.8 U χ U 10.8 nm^3^). The solvated system of protein was processed for energy minimization using the steepest algorithm up to a maximum 25,000 steps or until the maximum force (Fmax) is not greater than 1000 kJ/ mol nm which is the default threshold. The NVT and NPT ensembles for 50,000 steps (100 ps) at 300 K and 1 atm. Here, the system was firstly equilibrated using NVT ensemble followed by NPT ensemble. Then after the final Molecular Dynamic Simulation was performed for dock complex of [GVYFASTEK epitope docked against the HLA-A*11-01 allele (PDB ID: 5WJL)]. Finally, the simulations were evaluated RMSD and RMSF were calculated for complete episode of simulations. The all steps are kept similar except the final the molecular dynamics simulation was carried out for 50 ns long.

## 3. Results

### 3.1. Selection and Retrieval of Viral Protein Sequences

The *SARS-CoV-2,* strain was identified. Twelve viral protein sequences of *SARS-CoV2* against (Host: Human, Country: India) were retrieved from ViPR database and selected for the possible vaccine candidate identification (Table-1). These proteins were: Orf10 protein (**QIA98591.1)**, Orf8 protein **(QIA98589.1)**, Orf7a protein **(QIA98588.1)**, Orf6 protein **(QIA98587.1)**, Orf3a protein **(QIA98584.1)**, Membrane glycoprotein **(QIA98586.1)**, Envelope protein **(QIA98585.1)**, Surface glycoprotein **(QIA98583.1)**, Surface glycoprotein **(QHS34546.1)**, Nucleocapsid protein **(QII87776.1)**, Nucleocapsid protein **(QII87775.1)** and Nucleocapsidphosphoprotein **(QIA98590.1)**. The FASTA sequence of proteins mentioned in **(Supplementary File: S1)**

**Table 1:** SARS-CoV-2 (Host: Human, Country: India) viral protein sequence identification and retrieval via ViPR database (Viral pathogen database and analysis resource).

| *S.No.* | *Gene Symbol* | *Protein Name* | *GenBank Accession* | *GenBank Protein Accession* |
| --- | --- | --- | --- | --- |
| **1** | orf10 | Orf10 protein | MT050493 | QIA98591.1 |
| **2** | orf8 | Orf8 protein | MT050493 | QIA98589.1 |
| **3** | orf7a | Orf7a protein | MT050493 | QIA98588.1 |
| **4** | orf6 | Orf6 protein | MT050493 | QIA98587.1 |
| **5** | orf3a | Orf3a protein | MT050493 | QIA98584.1 |
| **6** | M | Membrane glycoprotein | MT050493 | QIA98586.1 |
| **7** | E | Envelope protein | MT050493 | QIA98585.1 |
| **8** | S | Surface glycoprotein | MT050493 | QIA98583.1 |
| **9** | S | Surface glycoprotein | MT012098 | QHS34546.1 |
| **10** | N | Nucleocapsid protein | MT163715 | QII87776.1 |
| **11** | N | Nucleocapsid protein | MT163714 | QII87775.1 |
| **12** | N | Nucleocapsid phosphoprotein | MT050493 | QIA98590.1 |

### 3.2. Physicochemical Property Analysis and Protein Antigenicity

The physicochemical property analysis like number of amino acids, the molecular weights, theoretical pI, extinction coefficients (in M-1 cm-1), Est. half-life (in mammalian cell), instability indexes, aliphatic indexes and grand average of hydropathicity (GRAVY) of the twelve proteins were predicted (Table-2). For antigenicity prediction threshold value kept at 0.4, all proteins were found to be antigenic (Table-3). The physicochemical study revealed that the surface glycoprotein **(QIA98583.1)** had the comparatively maximum extinction co-efficient of 148960M-1 cm-1 and lowest GRAVY value of −0.077. In addition, surface glycoprotein was stable and antigenic. We selected this surface glycoprotein for further analysis.

**Table 2:** Physiochemical property analysis of SARS-CoV-2 against (Host: Human, Country: India) viral proteins.

| Gene Symbol | Properties |  |  |  |  |  |  |  |
| --- | --- | --- | --- | --- | --- | --- | --- | --- |
|  | No. of amino acids | Molecular weight | Theoretical pI | Ext. coefficient (in M-1 cm-1) | Est. half-life (in mammalian cell) | Instability index | Aliphatic index | GRAVY (grand average of hydropathicity) |
| orf10 | 38 | 4449.23 | 7.93 | 4470 | 30 hours | 16.06 (stable) | 107.63 | 0.637 |
| orf8 | 121 | 13804.93 | 5.42 | 16305 | 30 hours | 46.24 (unstable) | 94.13 | 0.181 |
| orf7a | 121 | 13744.17 | 8.23 | 7825 | 30 hours | 48.66 (unstable) | 100.74 | 0.318 |
| orf6 | 61 | 7272.54 | 4.60 | 8480 | 30 hours | 31.16 (stable) | 130.98 | 0.233 |
| orf3a | 275 | 31122.94 | 5.55 | 58705 | 30 hours | 32.96 (stable) | 103.42 | 0.275 |
| M | 222 | 25146.62 | 9.51 | 52160 | 30 hours | 39.14 (stable) | 120.86 | 0.446 |
| E | 75 | 8365.04 | 8.57 | 6085 | 30 hours | 38.68 (stable) | 144.00 | 1.128 |
| S | 1273 | 141206.52 | 6.24 | 148960 | 30 hours | 33.01 (stable) | 84.82 | -0.077 |
| S | 1272 | 140972.27 | 6.16 | 147470 | 30 hours | 32.78 (stable) | 85.05 | -0.071 |
| N | 88 | 9827.08 | 10.23 | 8480 | 4.4 hours | 36.54 (stable) | 61.14 | -1.067 |
| N | 133 | 14363.88 | 11.37 | 8480 | 1 hours | 58.97 (unstable) | 44.21 | -1.170 |
| N | 419 | 45625.70 | 10.07 | 43890 | 30 hours | 55.09 (unstable) | 52.53 | -0.971 |

**Table 3:**
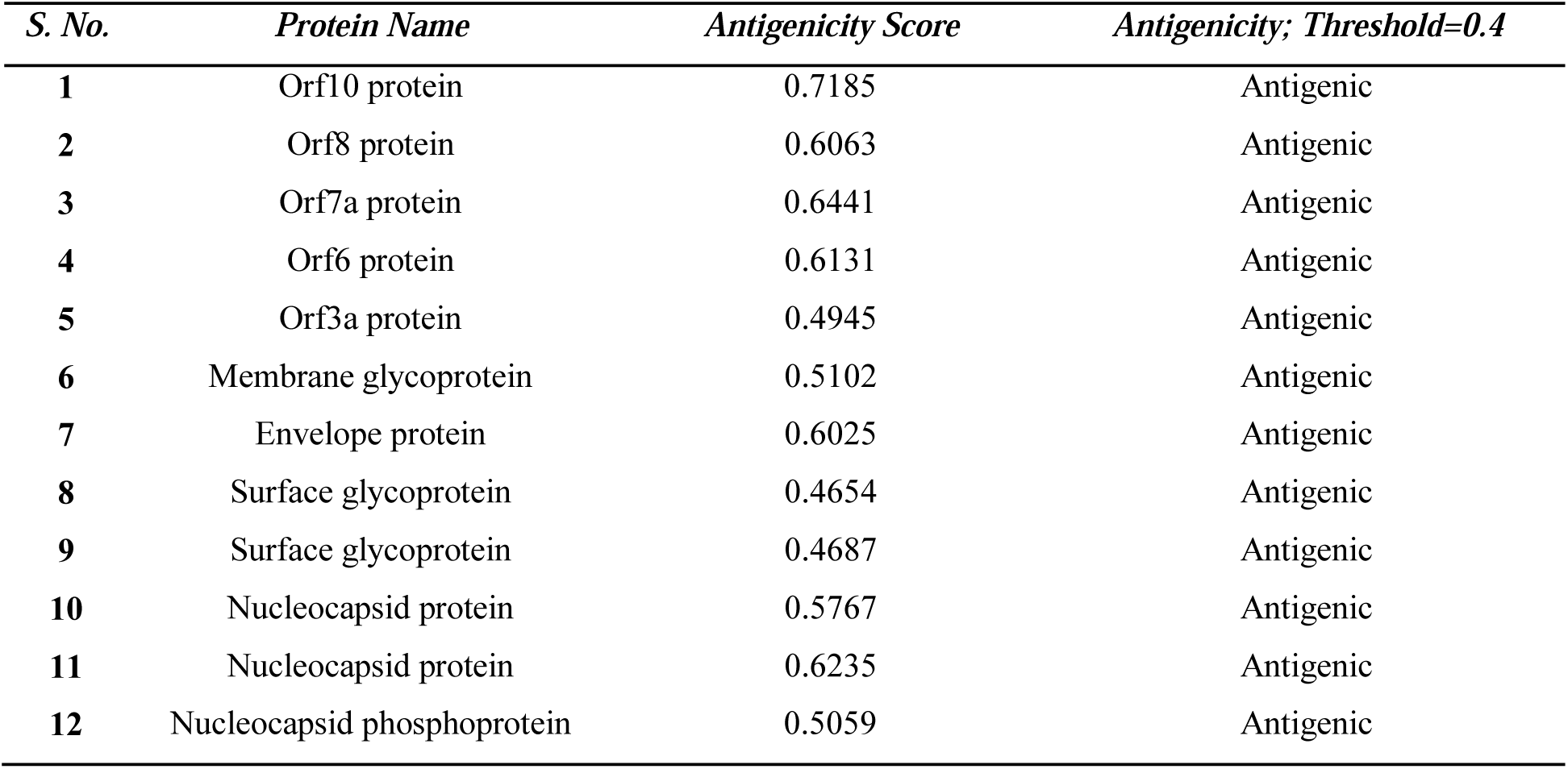
Antigenicity predication of SARS-CoV-2 viral proteins (Threshold value: 0.4)

### 3.3. T-cell and B-cell Epitope Prediction

The T-cell epitopes of MHC class-I was determined by NetMHCpan EL 4.0 prediction method of the IEDB server keeping the sequence length 9. The server generated epitopes further analyzed on the basis of the antigenicity scores (AS) and percentile scores, top ten potential epitopes were selected randomly for antigenicity, allergenicity, toxicity and conservancy tests. The server ranks the predicted epitopes based on the ascending order of percentile scores (Table-4a). The T-cell epitopes of MHC class-II (HLA DRB1*04-01 allele) of the protein was also determined by IEDB server (Table-4b), where the Sturniolo prediction methods was used. For protein, ten out of the top ranked epitopes were selected randomly for further analysis. Additionally, the B-cell epitopes of the protein was selected using Bipipered linear epitope prediction method of the IEDB server and the selection of epitopes were based on their higher lengths (Fig-1).

**Fig 1:**
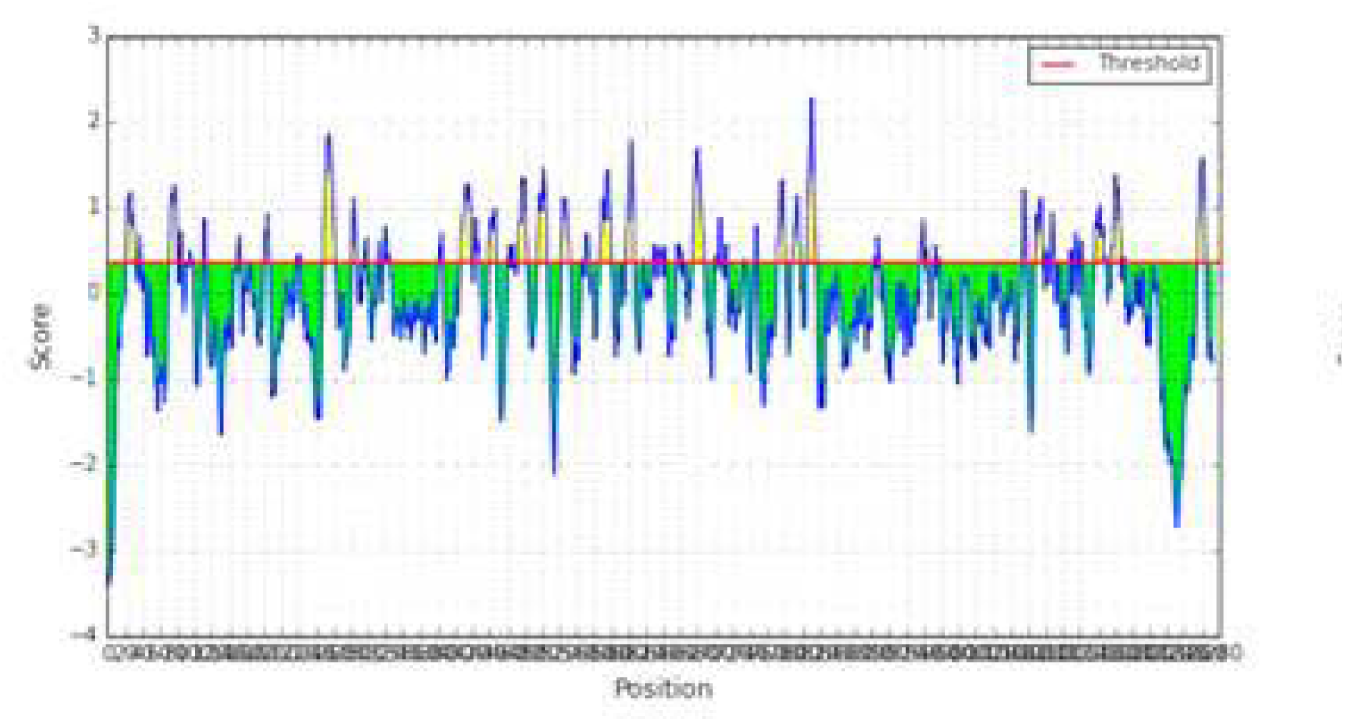
B cell Epitopes prediction: The graph of epitope prediction for surface glycoproteins QIA98583.1

**Table 4a:** MHC Class-I epitopes of Surface glycoprotein (QIA98583.1): Table represents topology, antigenicity, allergenicity, toxicity and conservancy analysis of protein.

| *Surface glycoprotein (QIA98583.1)* |  |  |  |  |  |  |  |  |  |
| --- | --- | --- | --- | --- | --- | --- | --- | --- | --- |
| Epitope | Start | End | Topology | Antigenicity | Score | Allergenicity | Toxicity | Minimum Identity | Conservancy |
| **GVYFASTEK** | 19 | 27 | Inside | Antigen | 0.7112 | Non-allergen | Non-toxic | 11.11% | 100% |
| **VTYVPAQEK** | 15 | 23 | Inside | Antigen | 0.8132 | Allergen | Non-toxic | 22.22% | 100% |
| **ASANLAATK** | 40 | 48 | Inside | Antigen | 0.7041 | Allergen | Non-toxic | 22.22% | 100% |
| **TLADAGFIK** | 57 | 65 | Inside | Antigen | 0.5781 | Non-allergen | Non-toxic | 22.22% | 100% |
| **TLKSFTVEK** | 22 | 30 | Inside | Non-antigen | 0.0809 | Allergen | Non-toxic | 11.11% | 100% |
| **NSASFSTFK** | 20 | 28 | Inside | Non-antigen | 0.1232 | Allergen | Non-toxic | 11.11% | 100% |
| **TEILPVSMTK** | 24 | 33 | Inside | Antigen | 1.4160 | Allergen | Non-toxic | 10.00% | 100% |
| **SSTASALGK** | 29 | 37 | Outside | Antigen | 0.6215 | Allergen | Non-toxic | 22.22% | 100% |
| **GTHWFVTQR** | 49 | 57 | Inside | Non-antigen | 0.0723 | Allergen | Non-toxic | 11.11% | 100% |
| **EILPVSMTK** | 25 | 33 | Inside | Antigen | 1.6842 | Allergen | Non-toxic | 11.11% | 100% |

**Table 4b:**
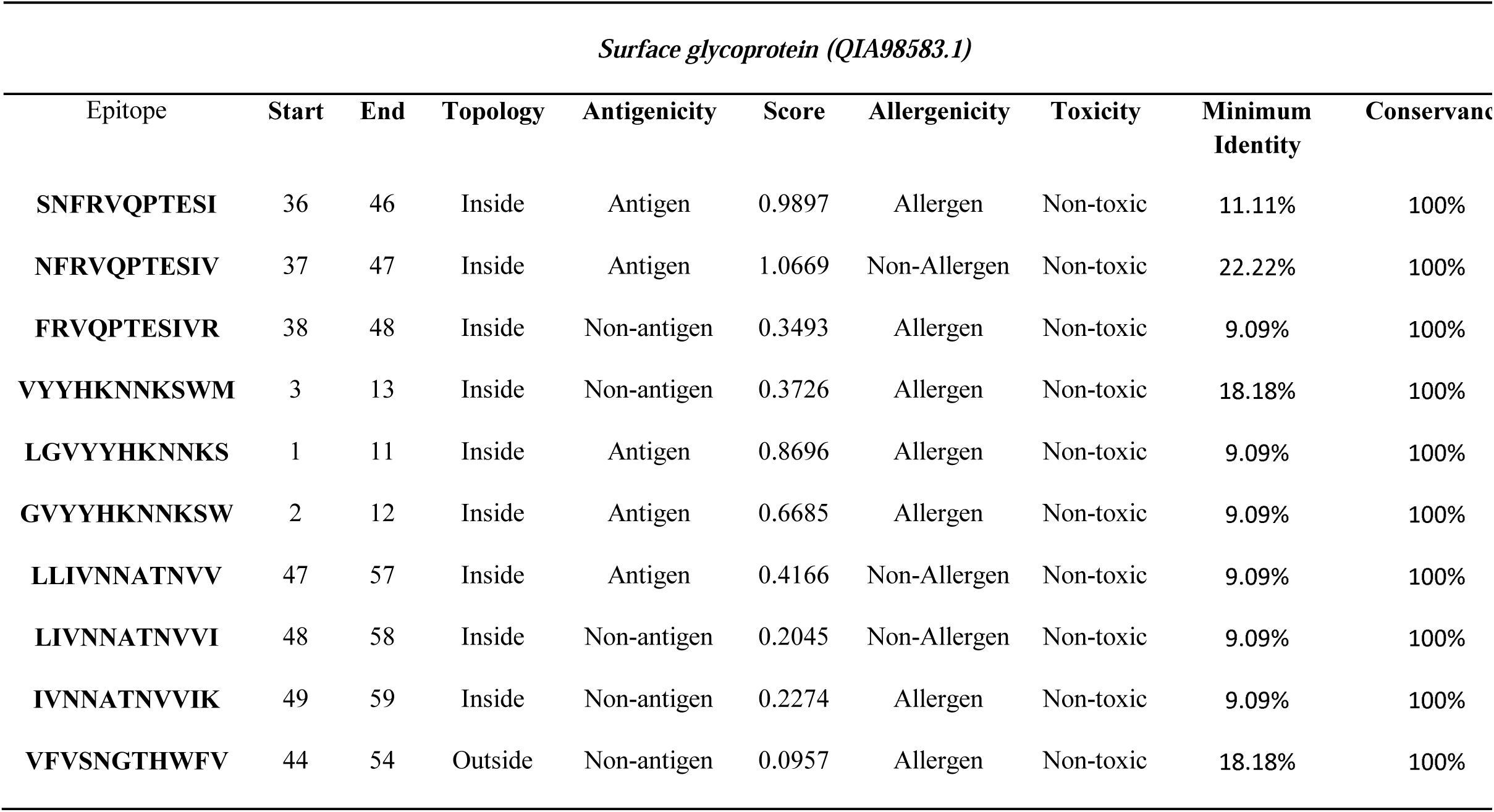
MHC Class II epitopes of Surface glycoprotein (QIA98583.1): Table represents topology, antigenicity, allergenicity toxicity and conservancy analysis of protein.

### 3.4. Topology Identification of the Epitopes

The topology of the selected epitopes was determined by TMHMM v2.0 server (http://www.cbs.dtu.dk/services/TMHMM/). The Table 4a and Table 4b represent the potential T-cell epitopes of selected surface glycoprotein. Table 5 shows the potential B-cell epitopes with their respective topologies.

**Table 5:** B cell epitopes of Surface glycoprotein (QIA98583.1): Table represents topology, antigenicity and allergenicity analysis of protein.

| *Surface glycoprotein (QIA98583.1)* |  |  |  |  |  |  |  |
| --- | --- | --- | --- | --- | --- | --- | --- |
| Epitope | Topology | Antigenicity | Allergenicity | Epitope | Topology | Antigenicity | Allergenicity |
| **RTQLPPAYTNS** | Inside | Antigen | Allergen | **QIAPGQTGKIAD** | Inside | Antigen | Non-Allergen |
| **SGTNGTKRFDN** | Inside | Antigen | Allergen | **YGFQPTNGVG YQ** | Outside | Antigen | Allergen |
| **LTPGDSSSGWTAG** | Outside | Antigen | Non-Allergen | **RDIADTTDAVRDPQ** | Inside | Antigen | Allergen |
| **VRQIAPGQTGKIAD** | Inside | Antigen | Non-Allergen | **QTQTNSPRRARSV** | Inside | Non-antigen | Non-Allergen |
| **YQAGSTPCNGV** | Inside | Non-antigen | Non-Allergen | **ILPDPSKPSKRS** | Outside | Antigen | Non-Allergen |

### 3.4. Antigenicity, allergenicity, toxicity and conservancy analysis of the epitopes

The selected T-cell epitopes found to be highly antigenic as well as non-allergenic, non-toxic, and had conservancy of greater than 90%. Among the ten selected MHC class-I epitopes and ten selected MHC class-II epitopes, total four epitopes were selected based on the mentioned criteria: GVYFASTEK, TLADAGFIK, NFRVQPTESI and LLIVNNATNV.

### 3.5. Cluster Analysis of the MHC Alleles

The cluster analysis of the MHC class-I alleles that will possibly interact with the predicted epitopes were carried out by online tool MHCcluster 2.0 (http://www.cbs.dtu.dk/services/MHCcluster/). The tool generates the clusters of the alleles in phylogenetic manner. The Results illustrate the outcome of the experiment where the red zone indicates strong interaction and the yellow zone corresponds to weaker interaction (Fig-2).

**Fig 2:**
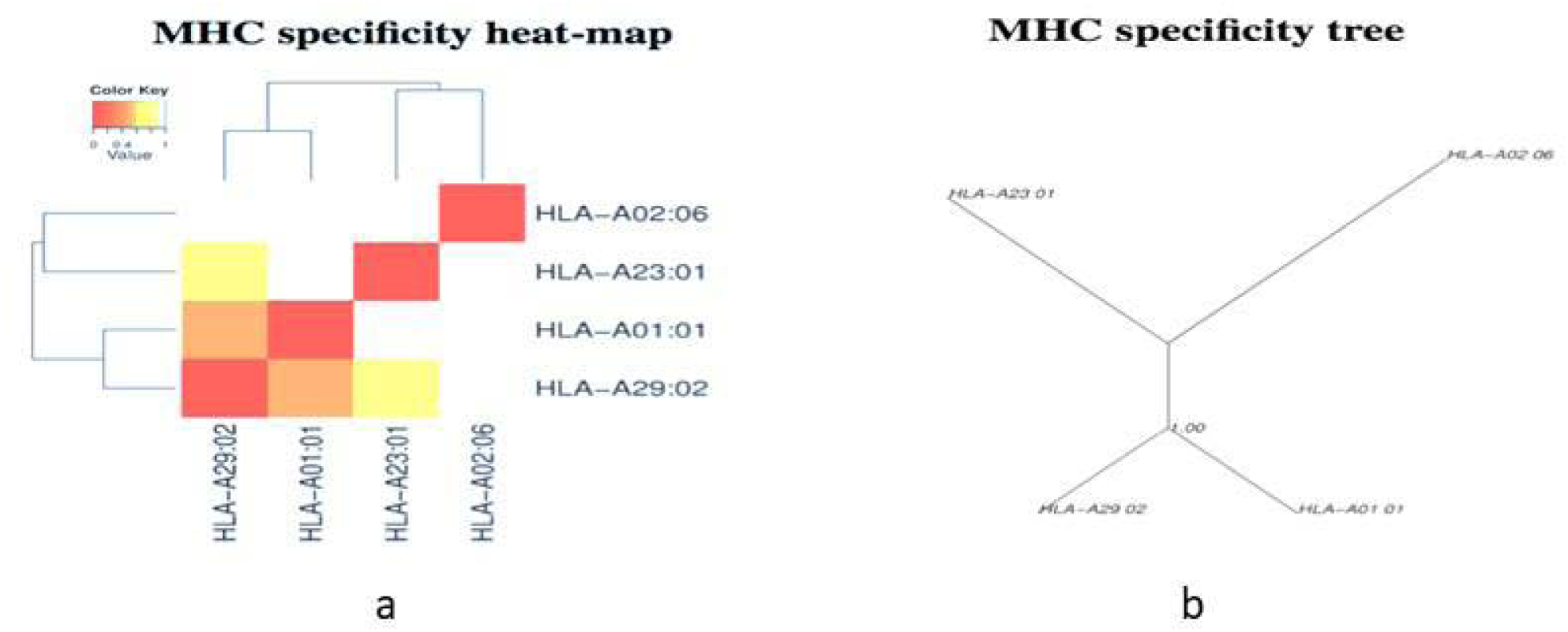
MHC Class Cluster Analysis (a) Heat map (b) Specificity tree Note: Here, red zone indicates strong interaction and the yellow zone corresponds to weaker interaction.

### 3.6. The 3D Structure prediction of the Epitopes

All the T-cell epitopes were subjected to 3D structure is predicted by the PEP-FOLD3 server. The predicted 3D structures were used for peptide-protein docking (Fig-3).

**Fig 3:**
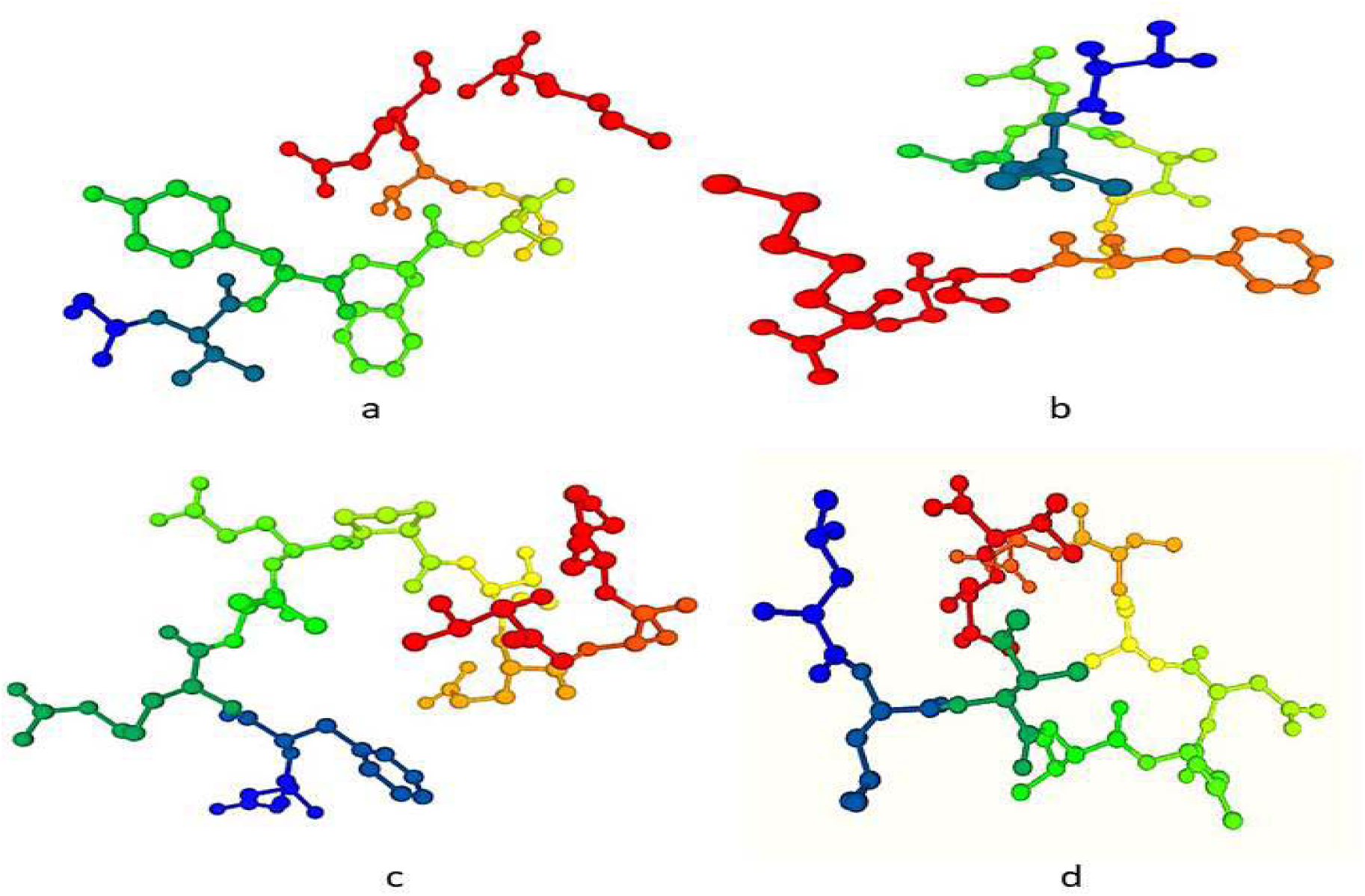
3D structure generation of T-cell epitopes by the PEP-FOLD3 server. Epitope representation: (a) GVYFASTEK (b) TLADAGFIK (c) NFRVQPTESIand (d) LLIVNNATNV.

### 3.7. Peptide-Protein Docking &Vaccine candidate’s prioritization

The Molecular Docking study was performed to observe, whether all the epitopes had the capacity to bind with the MHC class-I and MHC class-II molecule. The selected epitopes were docked with the HLA-A*11-01 allele (PDB ID: 5WJL) and HLA DRB1*04-01 (PDB ID: 5JLZ). The docking was performed using PatchDock online docking tool and then the results were refined by FireDock online server. Results were also analyzed by HPEPDOCK server **(Supplementary Figure: S2)**. Among the Four epitopes of selected glycoprotein QIA98583.1, GVYFASTEK (MHC class I epitope) showed the best result with the lowest global energy of **-**52.82. Further, docking pose was analyzed via Ligplot (Fig 4a) and docking site can be clearly visualized in (Fig 4b). We also identified highly antigenic and non-allergenic B-cell vaccine candidates LTPGDSSSGWTAG and VRQIAPGQTGKIAD from the selected Surface glycoprotein (QIA98583.1).

**Fig 4a:**
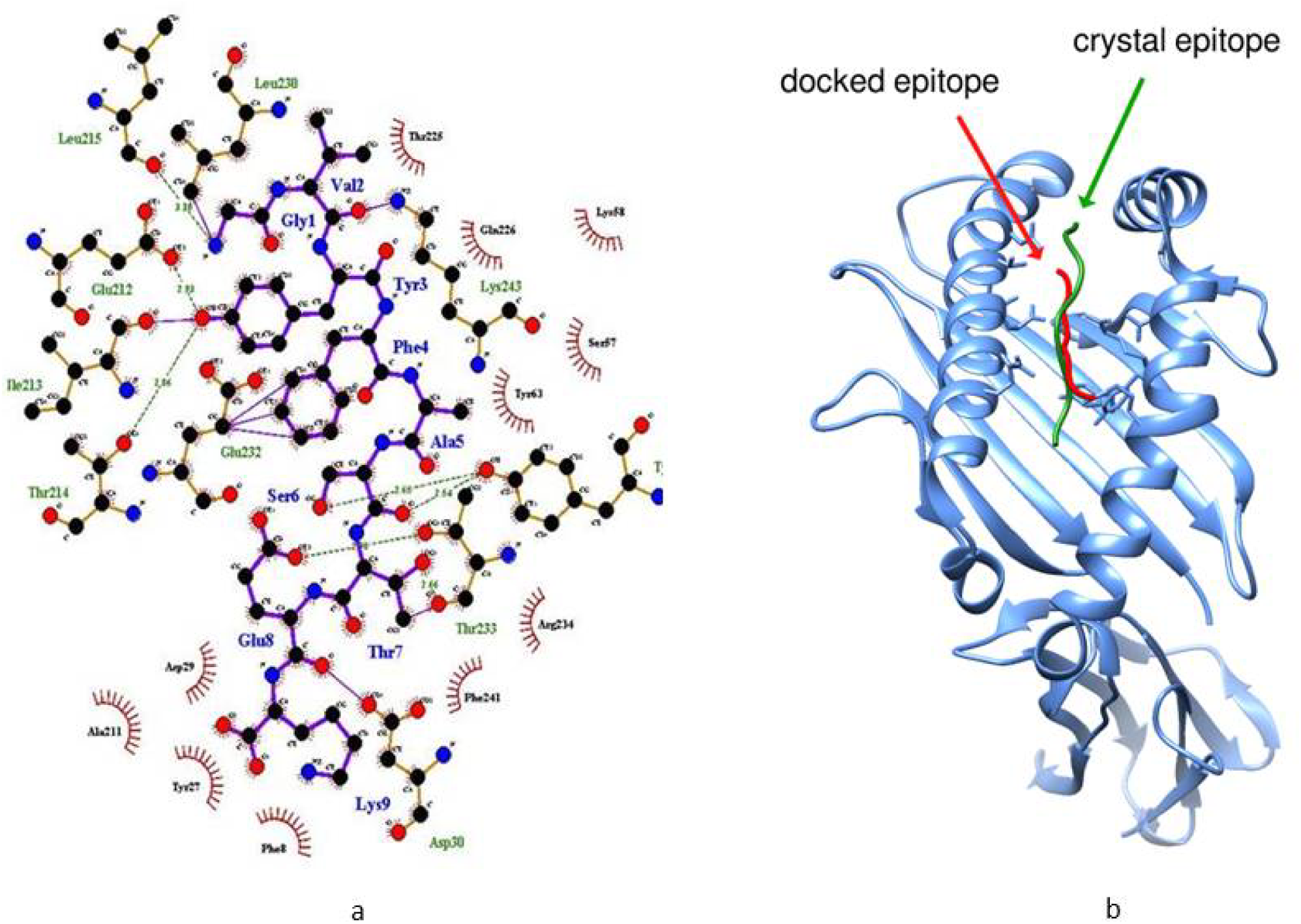
Docking pose analysis via LigPlot [GVYFASTEK epitope docking against the HLA-A *11-01 allele (PDB ID: 5WJL)] Note: Molecular Docking result showing protein -ligand interaction; where Oxygen (O), Nitrogen (N) and Carbon (C) atoms are represented in red, blue and black circle. Molecular docking analysis showing docking site of ligand (GVYFASTEK epitope) in our study is similar to ligand used in crystal structure of HLA-A*11-01 allele (PDB ID: 5WJL).

## Molecular Dynamics Simulation

Molecular Dynamics Simulation study of dock complex of GVYFASTEK epitope docked against the HLA-A*11-01 allele (PDB ID: 5WJL) was successfully executed for 50ns. The complex became stable throughout the simulation with RMSD fluctuation between 0.3-1.0 nm from the original position (Fig. 5a). In most cases, residues lying in the core protein regions have low RMSF while exposed loops have high RMSF values (Fig. 5b). As observed, the peaks in the graph show a value between 0.1 and 0.6 nm. Both these results indicate that the protein complexes were stable throughout MD simulations and thus proteins possess the ability to stability.

**Fig 5:**
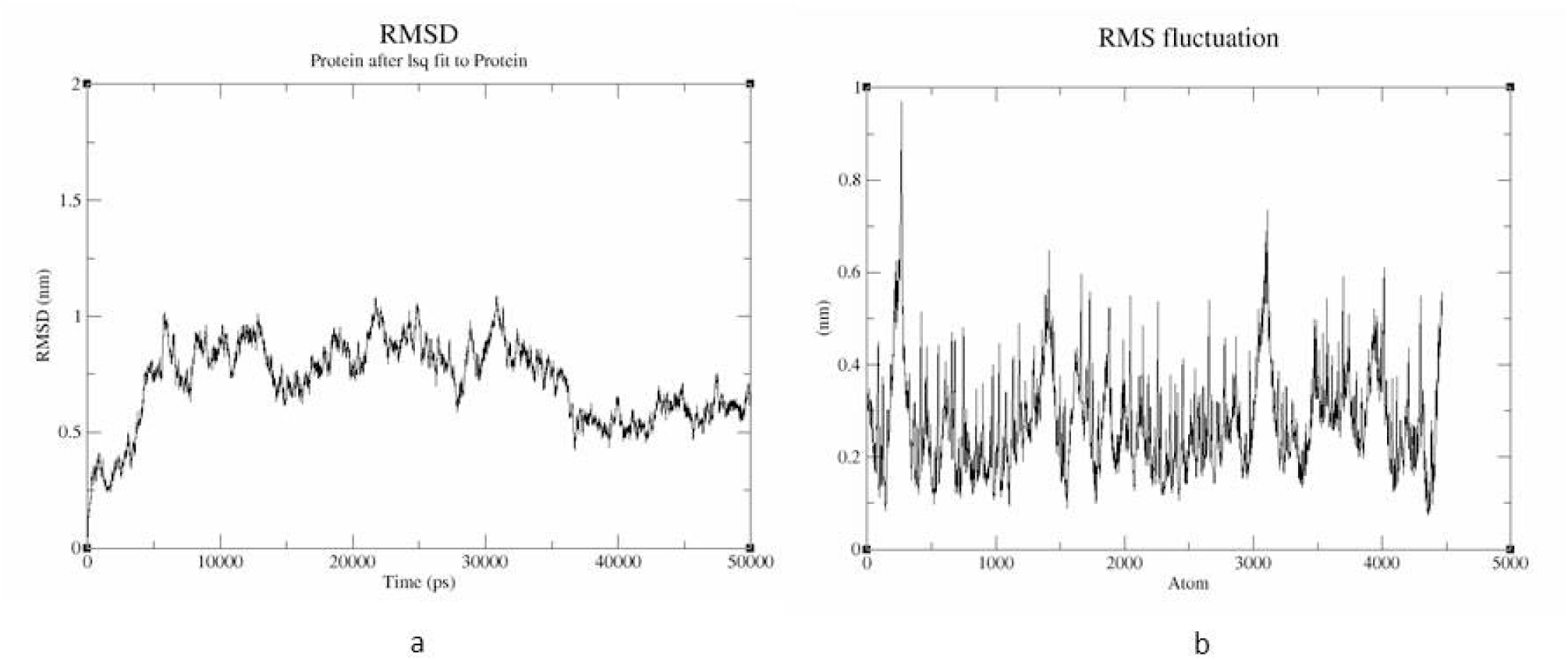
Molecular dynamics simulation-(a) RMSD graph (b) RMSF graph of dock complex [GVYFASTEK epitope docked against the HLA-A*11-01 allele (PDB ID: 5WJL)].

## 4. Discussion

Vaccine is enormously imperative and expansively formed therapeutic products. Millions of infants and people are getting vaccinated every year. However, the development and research processes of vaccines are expensive and occasionally, it requires countless months to prepare and advance an appropriate vaccine candidate towards killing of a pathogen. In today’s scenario, innumerable tools and approaches of immunoinformatics, Computer-aided drug design (CADD), bioinformatics and converse/reverse vaccinology stance extensively castoff towards the progression of vaccine preparations, which intern reduce prolong duration and price investment on vaccine expansion (Islam et al., 2020; María, Arturo, Alicia, Paulina, & Gerardo, 2017; Ullah et al., 2020).

In current study, the physicochemical study revealed that surface glycoprotein **QIA98583.1** exhibited upper most extinction co-efficient of 148960M-1 cm-1 and lowest GRAVY value of −0.077. In addition, this selected surface glycoprotein was highly stable (instability index of less than 40) and antigenic. The antigenicity of the protein was determined by VaxiJen V2.0 server. If a compound required variability index greater than 40, means the product is considered as unbalanced (Guruprasad, Reddy, & Pandit, 1990). The extinction coefficient referred to the quantity of light, that is captivated by a complex at a particular wavelength (Ikai, 1980; Pace, Vajdos, Fee, Grimsley, & Gray, 1995). Various physicochemical properties including the amount of amino acids, molecular mass/weight, theoretic pI, extinction co-efficient, uncertainty index, aliphatic index, GRAVY was resolute by ProtParam (https://web.expasy.org/protparam/) server.

In the immune system, two major functioning cells are B and T lymphocytic cells, which are responsible for several defended roles in the body. Once identified by an antigen presenting cell or APC (such as dendritic cell, macrophage etc.), the antigen is accessible by the MHC class-II molecule existing on the exterior of these usual APCs, to the helper T cell. Subsequently, the helper T cell comprehends CD4+ fragment on its exterior, it is correspondingly termed as CD4+ T cell. Later stimulated by APC, the helper-T cells subsequently stimulate the B cell and reasons to yield antibody constructing plasma B cell alongside memory B cell. The plasma B cell harvests many antibodies and the memory B cell purposes as the immunological, elongated tenure memory. Nevertheless, macrophage and CD8+ cytotoxic T cell are also triggered by the helper-T cell that abolishes the target antigen (Arpin et al., 1995; Cano & Lopera, 2013; Goerdt & Orfanos, 1999; Pavli, Hume, Van De Pol, & Doe, 1993; Tanchot & Rocha, 2003).

The possible B and T cell epitopes of the designated *SARS-CoV-2* viral protein was identified by the IEDB (https://www.iedb.org/) server. The IEDB server generates and ranks the epitopes on the basis of their antigenicity scores (AS) and percentile scores. The top ten MHC class-I and class-II epitopes were engaged for investigation. The topology of the precise epitopes was resolute by TMHMM v2.0 server (http://www.cbs.dtu.dk/services/TMHMM/). In all the inflammatory situations such as allergenicity, antigenicity, toxicity and conservancy examination, the T-cell epitopes those were found to be exceedingly antigenic with higher immune response without allergies, non-toxic, and required conservancy of over 90%. Amongst ten certain MHC class-I and ten selected MHC class-II epitopes of the protein, over-all four epitopes were designated based on the revealed measures: GVYFASTEK, TLADAGFIK, NFRVQPTESI and LLIVNNATNVV as well as antigenic and non-allergenic B-cell epitopes were selected for added vaccine candidate investigation. The cluster examination of the conceivable MHC class-I and MHC class-II alleles that might interrelate with the prophesied epitopes were performed by the online tool MHC cluster 2.0 (http://www.cbs.dtu.dk/services/MHCcluster/). The antigenicity, demarcated as the capability of a extraneous ingredient to act as an antigen and stimulate the B and T cell replies, over their epitope correspondingly baptized antigenic determinant portion (Fishman, Wiles, & Wood, 2015). The allergenicity is well-defined as the capability of that ingredient to act as an allergen and persuade latent allergic responses inside the host being (Andreae & Nowak-Wgrzyn, 2017).

Moreover, the cluster scrutiny of the MHC class-I and II alleles were similarly performed to categorize their association with each other and bunch them based on their functionality and foretold requisite specificity(Thomsen et al., 2013). Subsequently in the following steps, the peptide-protein docking was performed amongst the chosen epitopes and the MHC alleles. The MHC class-I epitopes stayed docked to the MHC class-I molecule (PDB ID: 5WJL) and the MHC class-II epitopes were docked with the MHC class-II molecule (PDB ID: 5JLZ) correspondingly. The peptide-protein docking was executed to evaluate the capability of the epitopes to quandary with their particular MHC molecules. Pre-docking was performed by UCSF-chimera and afterward, 3D structure generation of the epitopes was performed. The docking was executed by PatchDock and FireDock servers and analyzed by HPEPDOCK server constructed on global energy. The GVYFASTEK engendered the best scores in the peptide-protein docking respectively. All the vaccine candidates were proved to be potentially antigenic and non-allergenic, for this reason they should not cause any allergenic reaction within the host body. However, more *in vitro* and *in vivo* examinations should be performed to lastly approve the security, usefulness and potentiality of the predicted vaccines candidates.

## 5. Conclusion

In the face of enormous tragedy of suffering, demise and social adversity caused by COVID-19 pandemic. It is of the extremely importance to develop an effectual and safe vaccine against this highly pandemic disease. Bioinformatics, Reverse vaccinology and related technologies are widely used in vaccine design and development since these technologies reduce the cost and time. In this study, first the potential proteins belong to *SARS-CoV-2* against (host: human, country: India) are identified. Further, the potential B cell and T cell epitopes that can effectively elicit cellular mediated immune response related to these selected proteins were determined through robust processes. These potential T-cell epitope GVYFASTEK and B-cell epitopes LTPGDSSSGWTAG, VRQIAPGQTGKIAD, QIAPGQTGKIAD and ILPDPSKPSKRS play vigorous protagonist in the subunit and multi-epitope vaccine construction soon. In brief, reverse vaccinology vindicated as an authoritative means to recognize novel vaccine candidates and their consequential exact application. This study will lead to the research in an innovative and efficient direction and the outcome of our study will deliver a fast, reliable and significant platform in search of effective and timely cure of this treacherous pandemic disease, COVID-19 caused by *SARS-CoV2.*

## Acknowledgement

RKM acknowledges the financial support and award of Ramalingaswami fellowship from Department of Biotechnology, New Delhi, India. Authors highly acknowledge the Amity Institute of Biotechnology, Amity University Rajasthan, Jaipur and Dr. B. Lal Institute of Biotechnology, Jaipur.

## Compliance with Ethical Standards

## Ethical Approval

This article does not contain any studies with human participants or animals performed by any of the authors.

